# Kappa Opioid Receptor and Dynorphin Signaling in the Central Amygdala Regulates Alcohol Intake

**DOI:** 10.1101/663666

**Authors:** Daniel W. Bloodgood, Dipanwita Pati, Melanie M. Pina, Sofia Neira, J. Andrew Hardaway, Shivani Desai, Kristen M. Boyt, Richard D. Palmiter, Thomas L. Kash

## Abstract

Excessive alcohol drinking has been shown to modify brain circuitry to predispose individuals for future alcohol abuse. Previous studies have implicated the central nucleus of the amygdala (CeA) as an important site for mediating the somatic symptoms of withdrawal and for regulating alcohol intake. In addition, recent work has established a role for both the Kappa Opioid Receptor (KOR) and its endogenous ligand dynorphin in mediating these processes. However, it is unclear whether these effects are due to dynorphin or KOR arising from within the CeA itself or other input brain regions. To directly examine the role of preprodynorphin (PDYN) and KOR expression in CeA neurons, we performed region-specific conditional knockout of these genes and assessed the effects on the Drinking in the Dark (DID) and Intermittent Access (IA) paradigms. We then examined the effects of DID on PDYN and KOR modulation of CeA circuit physiology. Conditional gene knockout resulted in sex-specific responses wherein PDYN knockout decreased alcohol drinking in both male and female mice, whereas KOR knockout decreased drinking in males only. We also found that neither PDYN nor KOR knockout protected against anxiety caused by alcohol drinking. Lastly, a history of alcohol drinking did not alter synaptic transmission in PDYN neurons in the CeA of either sex, but excitability of PDYN neurons was increased in male mice only. Taken together, our findings indicate that PDYN and KOR signaling in the CeA plays an important role in regulating excessive alcohol consumption and highlight the need for future studies to examine how this is mediated through downstream effector regions.

## Introduction

Excessive alcohol drinking is a serious public health problem and was estimated to cost the United States over $249 billion in 2010 (Sacks, Gonzales, Bouchery, Tomedi, & Brewer, 2015). The majority of these expenses could be attributed to binge drinking, a form of alcohol consumption wherein individuals imbibe large amounts of alcohol within a short period of time and reach blood alcohol concentrations above 0.08% (80 mg/dL) (NIAAA). Individuals who engage in frequent binge drinking have a higher risk of later developing Alcohol Use Disorder (AUD) (Jennison, 2004; McCarty et al., 2004), and binge-alcohol consumption has been shown to engage the neural circuitry involved in the addiction cycle (Lowery-Gionta et al., 2012). Specifically, work from our lab has found that both the central nucleus of the amygdala (CeA) and the Bed Nucleus of the Stria Terminalis (BNST), components of the extended amygdala (Alheid & Heimer, 1988), are involved in binge-like alcohol consumption (Pleil, Rinker, et al., 2015; Rinker et al., 2017). These regions are important for regulating negative affective states, which are thought to underlie the negative reinforcing properties of alcohol (Koob & Le Moal, 2005).

Abundant within these brain regions is the kappa opioid receptor (KOR) and its endogenous ligand dynorphin (Lin et al., 2006; Mansour, Fox, Akil, & Watson, 1995). Activation of KOR results in dysphoria in human subjects (Pfeiffer, Brantl, Herz, & Emrich, 1986), and recently it has been shown that there is reduced KOR occupancy in the amygdala of patients with an AUD (Vijay et al., 2018). In animal models, systemic treatment with KOR antagonists has been shown to reduce ethanol self-administration (Walker & Koob, 2007), stress-enhanced drinking (Anderson, Lopez, & Becker, 2016), and ethanol dependence-induced increases in anxiety (Valdez & Harshberger, 2012). Additionally site-specific KOR antagonism in the CeA decreases ethanol self-administration in dependent animals (Kissler et al., 2014) and alcohol consumption in a model of binge-like drinking (Anderson et al., 2018). Anderson et al. (2018) also found that chemogenetic inhibition of preprodynorphin (PDYN) neurons in the CeA reduced binge-like drinking, suggesting that the source of dynorphin release may be within the CeA. However, one limitation of these experiments is that PDYN neurons co-express a variety of other neuropeptides such as corticotropin releasing factor, somatostatin, and neurotensin (Kim, Zhang, Muralidhar, LeBlanc, & Tonegawa, 2017). Additionally, as these neurons form many local inhibitory connections (Pomrenze et al., 2015), it is unclear whether the results of the chemogenetic experiment are due directly to release of dynorphin, GABA or some other co-expressed neuropeptide.

To examine more closely how this system may work to regulate ethanol consumption, we performed dual *in situ* hybridization of *Pdyn* and *Oprk1* mRNA (the *Oprk1* gene encodes KOR protein) and found that they were expressed in separate, relatively non-overlapping neuronal populations in the CeA. We then separately knocked out *Pdyn* and *Oprk1* and found that these manipulations resulted in reductions in alcohol drinking in a sex-specific manner. Finally, we performed slice electrophysiology recordings from mice that underwent binge-like drinking and found that a history of alcohol drinking resulted in sex-specific alterations in PDYN neuron excitability with no effect on synaptic transmission.

## Materials and Methods

### Subjects

Experiments were performed on adult male and female mice between 8-10 weeks of age at the beginning of the experiments. *Pdyn^IRES-Cre^* and *Gt(ROSA26)Sor^loxSTOPlox-L10-GFP^* (Krashes et al., 2014), *Oprk1^lox/lox^* conditional-knockout mice (Chefer, Bäckman, Gigante, & Shippenberg, 2013) were generated as described and bred in house at UNC. Additionally, *Pdyn^lox/lox^* mice were generated as described below and were subsequently transferred and bred in house at UNC. All mice were group housed in colony rooms on a 12:12 hr light-dark cycle (lights on at 7 a.m.) with *ad libitum* access to rodent chow and water. All animal procedures were performed in accordance with the Institutional Animal Care and Use Committee at the University of North Carolina at Chapel Hill.

### Generation and Validation of Pdyn^Lox^ Conditional Knockout Mice

The 5’ arm of the targeting construct (3.8 kb) was PCR amplified from a C57Bl/6 BAC clone using Q5 DNA polymerase with *Pac*1 site at the 5’ end and *Xba*1 site at the 3’ end (Supplementary Figure 2a). A synthetic loxP site was inserted into the *BamH*1 site 5’ of the first coding exon. The 3’ arm of the targeting construct (3.05 kb) was PCR amplified with *Sal*1 site at the 5’ end and *Not*1 site at the 3’ end. The two arms were cloned into the polylinkers of a targeting vector with a loxP site, a frt-flanked SV40-neomycin resistance gene for positive selection, *Pgk*-DTa and HSV-TK genes for negative selection. The targeting construct was electroporated into G4 ES cells and correct targeting was established by Southern blot of DNA digested with *Bcl*1 using a 32P-labelled probe located beyond the 3’ arm of the targeting construct. Seven of 60 clones were correctly targeted and all of them retained the distal loxP site assessed using PCR primers (5’ GACTCACTTGTTTGCTGGAGAG and 5’ CAGAGTACGTGGATTGTCACAG) flanking the distal loxP site. Several clones were injected into blastocysts of C57Bl/6J mice and then transferred to pseudo-pregnant females. One of the clones that gave high percentage of chimeric mice, was bred with mice expressing FLP recombinase to remove the NeoR gene. The mice were then continuously backcrossed to C57BL/6 mice. Routine genotyping was performed using the primers indicated above.

### Stereotaxic Surgeries

Mice were anesthetized with 4% isoflurane and placed in a stereotaxic frame (Kopf Instruments, Tujunga, CA, USA) and maintained under 1-2% isoflurane for the duration of the surgery. The skull was exposed and burr holes were made with a drill over the coordinates of the Central Amygdala (M/L +/- 2.90, A/P −1.20, D/V −4.70). A Hamilton Syringe (Hamilton Company, Reno, NV) was used to deliver 200 nL of an adeno associated virus encoding cre recombinase with a translationally fused green fluorescent protein (AAV5-*Camk2a*-Cre:GFP) or control vector (AAV5-*Camk2a*-eGFP) into the CeA. The fused Cre:GFP variant is nuclear-restricted resulting in a more punctate pattern of labeling compared to the cytosolic GFP control vector despite being matched in terms of injection volume. The virus was infused at a rate of 0.1 μL per minute and left in place for at least 5 minutes to ensure diffusion.

### Histology

Following the completion of behavioral experiments, mice were deeply anesthetized with an overdose of tribromoethanol (250 mg/kg, intraperitoneal) and transcardially perfused with 0.1 M phosphate buffered saline (4°C, pH 7.4) followed by 4% paraformaldehyde in PBS. Brains were post-fixed for 24 hours in 4% PFA and then cryoprotected in 30% sucrose in PBS. Coronal sections containing the CeA were cut at a thickness of 45 μm using a Vibratome (Leica VT1000S, Buffalo Grove, IL, USA) and stored in a 50% glycerol/PBS solution.

### In Situ Hybridization

Mice were anesthetized with isoflurane and rapidly decapitated. Brains were dissected and flash frozen on dry ice for 15 min and stored at −80°C until sectioned for in situ hybridization (ISH). Brain slices (16 µm) containing the CeA were obtained on a Leica CM 3050S cryostat (Leica Biosystems, Nussloch, Germany) at −20°C and were mounted directly onto microscope slides. Slides were stored at −80°C until tissue was processed for ISH. RNAscope ISH was conducted using the Multiplex Fluorescence Assay following the manufacturer’s protocol (Advanced Cell Diagnostics, Newark, CA, USA). The probes used to assess gene expression were purchased from Advanced Cell Diagnostics and are as follows: PDYN-C1 (Accession number NM_018863.3), OPRK1-C2 (Accession Number NM_001204371.1), and EGFP-C2 (Accession Number U55763.1). Slides were coverslipped with ProLong Gold Mounting Medium containing DAPI (Fisher Scientific, Pittsburgh, PA, USA) and stored at 4°C until they were imaged.

### Microscopy

Correct placements for viral infusions were verified using a wide-field epifluorescent microscope (BX-43, Olympus, Waltham, MA, USA) using a stereotaxic atlas (Franklin & Paxinos, 2008). Images for the quantification of fluorescent in situ hybridization were collected on a Zeiss 800 Laser Scanning Confocal Microscope (20x objective, NA=0.8) (Carl Zeiss, Jena, Germany). Regions of Interest (ROIs) containing the Central Lateral (CeL) and Central Medial (CeM) were annotated using Zeiss Zen 2 Blue Edition software. Manual quantification of PDYN positive and KOR positive cells was performed using manual counts with the cell counter plugin in FIJI (Curtis Rueden, LOCI, University of Wisconsin-Madison, Madison, WI, USA).

### qPCR on Tissue Punches

Mice were anesthetized with isoflurane and rapidly decapitated. The brain was dissected out and 1-mm coronal sections were made using a brain block. Slices were flash frozen on dry ice and tissue punches containing bilateral samples from the CeA were taken and stored at −80°C until future use. Total RNA from tissue punches was extracted using Direct-zol RNA miniprep kit (Zymo Research, Irvine, CA, USA). Reverse transcription and qPCR were performed using Superscript II and Applied Biosystems Taqman Assay on a StepOnePlus Real Time PCR System (Thermo Fisher Scientific, Waltham, MA, USA). The following TaqMan assay probes were purchased from Invitrogen (USA): ACTB Mm00607939_s1, *Oprk1* Mm01230885_m1, *Pdyn* Mm00457573_m1. Three technical replicates were averaged together to compute mean cycle threshold (Ct) values for the housekeeping gene and experimental genes of interest. The Ct value for β-actin was then subtracted from the Ct values for *Oprk1* and *Pdyn* to compute normalized ΔCt values for each sample.

### Alcohol and Tastant Drinking Procedures

All alcohol and tastant drinking experiments were conducted with mice single housed in a reverse light cycle space (lights off at 7am, lights on at 7pm). For the duration of all experiments, mice were maintained on an Isopro RMH 3000 (LabDiet, St. Louis, MO, USA) diet, as this been shown to result in the highest levels of ethanol consumption (Marshall et al., 2015). All animals were transferred to the reverse light space to acclimate to the chow and light cycle for at least one week prior to the start of experiments.

Drinking in the Dark (DID) was carried out as described (Lowery-Gionta et al., 2012; Pleil, Rinker, et al., 2015; Rhodes, Best, Belknap, Finn, & Crabbe, 2005). Briefly, on days 1–3 beginning 3 hr into the dark cycle, water bottles were removed from all cages and replaced with a bottle containing 20% (v/v) ethanol solution. Mice had 2 hr of access to ethanol, after which the ethanol bottles were removed from cages and water bottles were replaced. The same procedure was followed on day 4 except that ethanol access was extended to 4 hr. Bottle weights were recorded after 2 and 4 hr of access to ethanol on day 4. Additionally, a bottle was placed on an empty cage during all experiment days in order to record the amount of volume loss due to ethanol drip alone. This value was generally less than 0.1 mL/2 hr and was subtracted from the raw daily intake. The difference in start and end bottles weights minus the drip value was corrected for the density and concentration of ethanol and divided by the mouse’s body weight to obtain normalized intake g/Kg. Days 1-4 comprised one cycle of DID which was repeated for a total of 4 weeks, a point at which we have previously observed dynorphin and KOR modulation of binge-like drinking in the CeA (Anderson et al., 2018).

Following completion of DID experiments, cage tops were replaced with lids designed for 2-bottle drinking. Following 3 days of acclimation to bottles containing only water, animals began Intermittent Access to Ethanol (IA) as described previously (Hwa et al., 2011). Briefly, on Monday, Wednesday, and Friday mornings 3 hr into the dark cycle, one of the water bottles was replaced with 20% ethanol (w/v). Animals then had access to ethanol for 24 hr at which point the difference in weights of the water and ethanol bottles was measured. Drip values for water and ethanol bottles were calculated separately and used to calculate normalized consumption using the method described above. The side of the ethanol bottle (left or right) was counterbalanced across session to prevent the development of a side preference. Animals underwent IA for 2 wk to assess the effects of the genetic knockout on total fluid consumption and alcohol preference.

Ethanol-naïve animals underwent drinking of aversive or palatable solutions using the same procedures as the drinkers. However, the animals always had access to one bottle containing water and the other containing the tastant solution of interest. The concentrations of quinine (0.3, 0.6, and 0.9 mM), saccharin (0.33% and 0.66%), and sucrose (1%) were chosen based on values previously used to characterize global PDYN and KOR knockout mice (Blednov, Walker, Martinez, & Harris, 2006; Kovacs et al., 2005). The bottles for the tastant solution were changed 3 hr into the dark cycle and the animals had access to each solution for 24 hr thereafter. The weight of the tastant bottles and the drip bottles were recorded the following day and replaced on the opposite side as the previous day. The difference in consumption was averaged across the two days to obtain a single value for the preference of each solution. The bottle choice assay began with the lowest concentration of tastant in the series (0.3 mM quinine or 0.33% saccharin), which was then increased over the course of the next 6 days. This was then followed by 2 days of washout during which only water was available. We then began the preference test with the lowest concentration of the next series of solutions. The order of presentation of each solution type (palatable or aversive) was counterbalanced across cohorts within an experiment.

### Elevated Plus Maze

The Elevated Plus Maze (EPM) (Med Associates, St. Albans, VT) was made of white-and-black plastic and consisted of two open arms (75 × 7 cm) and two closed arms (75 × 7 × 25 cm) adjoined by a central area (7 × 7 × 25 cm). The arms were arranged in a plus configuration with arms of the same type (open or closed) opposite of each other. The maze was elevated 75 cm with light levels maintained at 15 lux throughout the experiment. Mice were placed in the center of the EPM and allowed to explore freely. The EPM was cleaned with 70% ethanol between each trial. Movements were video recorded and analyzed using Ethovision 9.0 (Noldus Information Technologies). The primary measures of reduced anxiety-like behavior were time spent in the open arm and number of entries into the open arm. One mouse in the PDYN knockout naïve male mice was excluded from the analysis as the subject exhibited only one entry into the closed arm. EPM testing was conducted 8 hrs after the last DID binge session (approximately 10pm or 3 hr in the light cycle). Testing in ethanol naïve animals was performed at the same time of day with respect to the light/dark cycle.

### Slice Electrophysiology

We performed whole-cell electrophysiology experiments similar to those published previously (Crowley et al., 2016; Lowery-Gionta et al., 2012). Briefly, 300-µm coronal slices containing the CeA were prepared on a vibratome (Leica VT1200, Leica, Wetzlar, Germany) from mice rapidly decapitated under isoflurane. The brains were removed and placed in ice-cold modified high sucrose artificial cerebrospinal fluid (aCSF) containing the following (in mM): 194 sucrose, 20 NaCl, 4.4 KCl, 2 CaCl_2_, 1 MgCl_2_, 1.2 NaH_2_PO_4_, 10.0 glucose, and 26.0 NaHCO_3_. Slices were then transferred to normal aCSF maintained at approximately 30°C (Warner Instruments, Hamden, Connecticut) containing the following (in mM): 124 NaCl, 4.4 KCl, 2 CaCl_2_, 1.2 MgSO_4_, 1 NaH_2_PO_4_, 10.0 glucose, and 26.0 NaHCO_3_. Slices were placed in a holding chamber where they were allowed to rest for at least one hour. Slices were continuously bubbled with a 95% O_2_ / 5% CO_2_ mixture throughout slicing and experiments. Thin-walled borosilicate glass capillary recording electrodes (3–6 MΩ) were pulled on a Flaming-Brown micropipette puller (Sutter Instruments, Novato, CA).

Following rupture of the cell membrane, cells were allowed to rest and equilibrate to the intracellular recording solutions (below). Signals were acquired via a Multiclamp 700B amplifier (Molecular Devices, Sunnyvale, California), digitized at 10 kHz and filtered at 3 kHz. Current clamp experiments were analyzed using Clampfit 10.2 software (Molecular Devices) and spontaneous synaptic transmission experiments were analyzed using Mini Analysis version 6.0.7 (Synaptosoft, Decatur, GA). Access resistance was monitored continuously throughout the experiment, and when it deviated by more than 20% the experiment was discarded. No more than two cells per animal were included in each experiment. For current clamp experiments, cells were recorded using a potassium-gluconate based internal recording solution containing the following (in mM): 135 K-gluc, 5 NaCl, 2 MgCl_2_, 10 HEPES, 0.6 EGTA, 4 Na_2_ATP, 0.4 Na_2_GTP. Experiments were conducted both at resting membrane potential (RMP) and −70mV.

Lidocaine N-ethyl bromide (1 mg/mL) was included in the intracellular recording solution to prevent postsynaptic sodium spikes for all voltage-clamp experiments. For voltage-clamp experiments requiring the simultaneous recording of excitatory and inhibitory events within the same neuron, a cesium-methanesulfonate based intracellular recording solution containing the following (in mM) was used: 135 Cs-meth, 10 KCl, 1 MgCl_2_, 0.2 EGTA, 4 MgATP, 0.3 GTP, 20 phosphocreatine. Excitatory events were recorded at −55mV and inhibitory events were recorded at +10mV. E/I synaptic drive ratio was calculated using the following formula: (EPSC frequency X amplitude)/ (IPSC frequency X amplitude).

For experiments examining the effects of KOR modulation of GABAergic transmission, a high chloride Cs-based internal solution was used (in mM): 117 Cs-Gluc, 20 HEPES, 0.4 EGTA, 5 TEA, 2 MgCl_2_, 4 Na_2_ATP, 0.4 Na_2_GTP. Electrode stimulation-evoked GABAergic inhibitory postsynaptic currents (eIPSCs) were pharmacologically isolated using 3mM kynurenic acid to block α-amino-3-hydroxy-5-methyl-4-isoxazole-propionic acid and N-methyl-D-aspartate receptor-dependent postsynaptic currents. Evoked IPSCs were induced via twisted bipolar nichrome wire placed dorsal to the recording electrode, 100 μm to 500 μm dorsal from the recorded neuron. Electric output was set to stimulate at 0.1 Hz, between 2 V and 16 V with a 100 microsecond to 150 microsecond duration. Following 5 minutes of a stable baseline, 1 μM U-69593 was applied for 10 minutes followed by 5 minutes of washout. Evoked IPSC experiments were analyzed by measuring the average peak amplitude of the synaptic response per minute, which was normalized to the baseline period 5 minutes immediately preceding application of the drug.

### Data Analysis and Statistics

Data are displayed as means ± SEM. For changes in the proportion of overlapping PDYN and KOR neurons after alcohol drinking, groups were compared with a χ^2^ test. Changes in mRNA expression were compared with a two-sample *t* test with Welch’s correction. Alcohol consumption and preference were analyzed as within subjects repeated measures ANOVA. Measures of anxiety-like behavior in the elevated plus maze were compared with a two-way (alcohol treatment X gene knockout) factorial ANOVA. Changes in synaptic transmission and excitability were assessed with either a repeated measures ANOVA or a two-sample *t* test with Welch’s correction where appropriate. All analyses were performed with R statistical software version 3.3.2.

## Results

### PDYN- and KOR-Expressing Neurons Form Separate Populations in the CeA and The Expression of Which is Not Altered Following Alcohol Drinking

Many studies have implicated the CeA as an important locus for regulating both dependent and non-dependent forms of alcohol drinking (Funk, O’Dell, Crawford, & Koob, 2006; Lowery-Gionta et al., 2012; Sparrow et al., 2012). Indeed, it has been shown that antagonizing KOR (Kissler et al., 2014) and chemogenetically inhibiting PDYN neurons (Anderson et al., 2018) can both decrease alcohol consumption. Despite evidence showing that both PDYN and KOR are expressed in the CeA, the circuit mechanism underlying this effect remains unclear. To assess the expression of PDYN and KOR, and how it may be affected by a history of alcohol drinking, we exposed C57BL/6J mice to 3 cycles of Drinking in the Dark (DID). Eight hours after the final binge session, animals were sacrificed for *in situ* hybridization and qPCR experiments (Figure 1a). Interestingly, we found that *Pdyn* and *Oprk1* mRNA were expressed on relatively distinct populations of neurons (Degree of co-localization: Lateral subdivision (CeL) *Mean*=22.8% ± 3.9%; Medial subdivision (CeM) *Mean*=18.5% ± 2.6%), and that the proportion of PDYN+, KOR+, and colocalized neurons was unaltered by ethanol drinking (Figure 1b-c). Additionally, *Pdyn* and *Oprk1* mRNA levels were unchanged following DID (Figure 1d-e). These findings suggest that although PDYN and KOR may be involved in the regulation of alcohol consumption, their gene expression levels are unaffected by a history of alcohol drinking.

**Figure 1.**
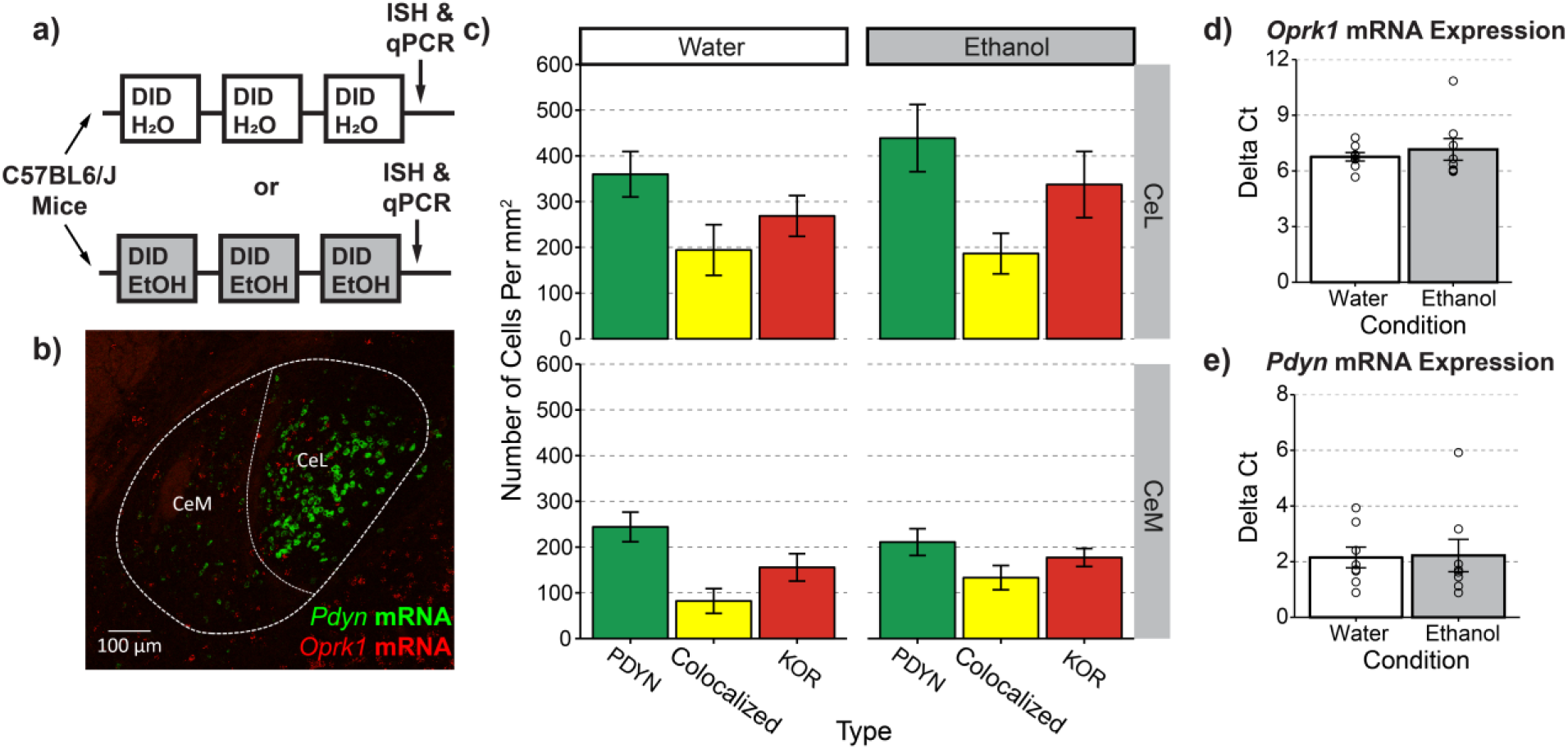
PDYN and KOR expression in CeA neurons is unaltered by a history of ethanol drinking. (a) Timeline for experimental procedures. (b) Representative image of *Pdyn* and *Oprk1* mRNA expression in the CeA (c) Quantification of PDYN+ and KOR+ cell counts following experimental manipulations. Ethanol treatment did not affect the proportion of PDYN+, KOR+, and colocalized neurons Χ^2^(2)=1.494, p=0.474 (d) Ethanol treatment did not affect *Oprk1* gene expression t(9.04)=0.64, p= 0.534 or (e) *Pdyn* gene expression t(11.899)=0.108, p=0.916. (c) n=12 images from N=3 ethanol or water drinking mice (d-e) punches from n=8 water and n=8 ethanol drinking animals

### Knockout of KOR in CeA Reduces Alcohol Consumption and Preference in a Sex-Specific Manner

Previous studies employing pharmacological approaches have shown that KOR antagonism in the CeA can decrease alcohol intake (Anderson et al., 2018; Kissler et al., 2014). However, it remains unclear whether this is mediated by KOR expression on CeA neurons or on presynaptic terminals from other brain regions. To establish a causal role for KOR signaling in CeA neurons, we performed a conditional knockout of KOR in young adult male and female mice. *Oprk1^lox/lox^* mice (Chefer et al., 2013) were infused with an AAV encoding Cre recombinase fused to eGFP or eGFP only as a control (Figure 2a-c). Alcohol drinking experiments began three weeks after virus injection, a time point we have previously demonstrated is sufficient for AAV-Cre driven knockout of KOR (Crowley et al., 2016). As predicted by the findings of Anderson et al. (2018), CeA KOR knockout resulted in a significant reduction in the amount of alcohol consumed across 4 wk of DID in male mice (Figure 2d-f).

**Figure 2.**
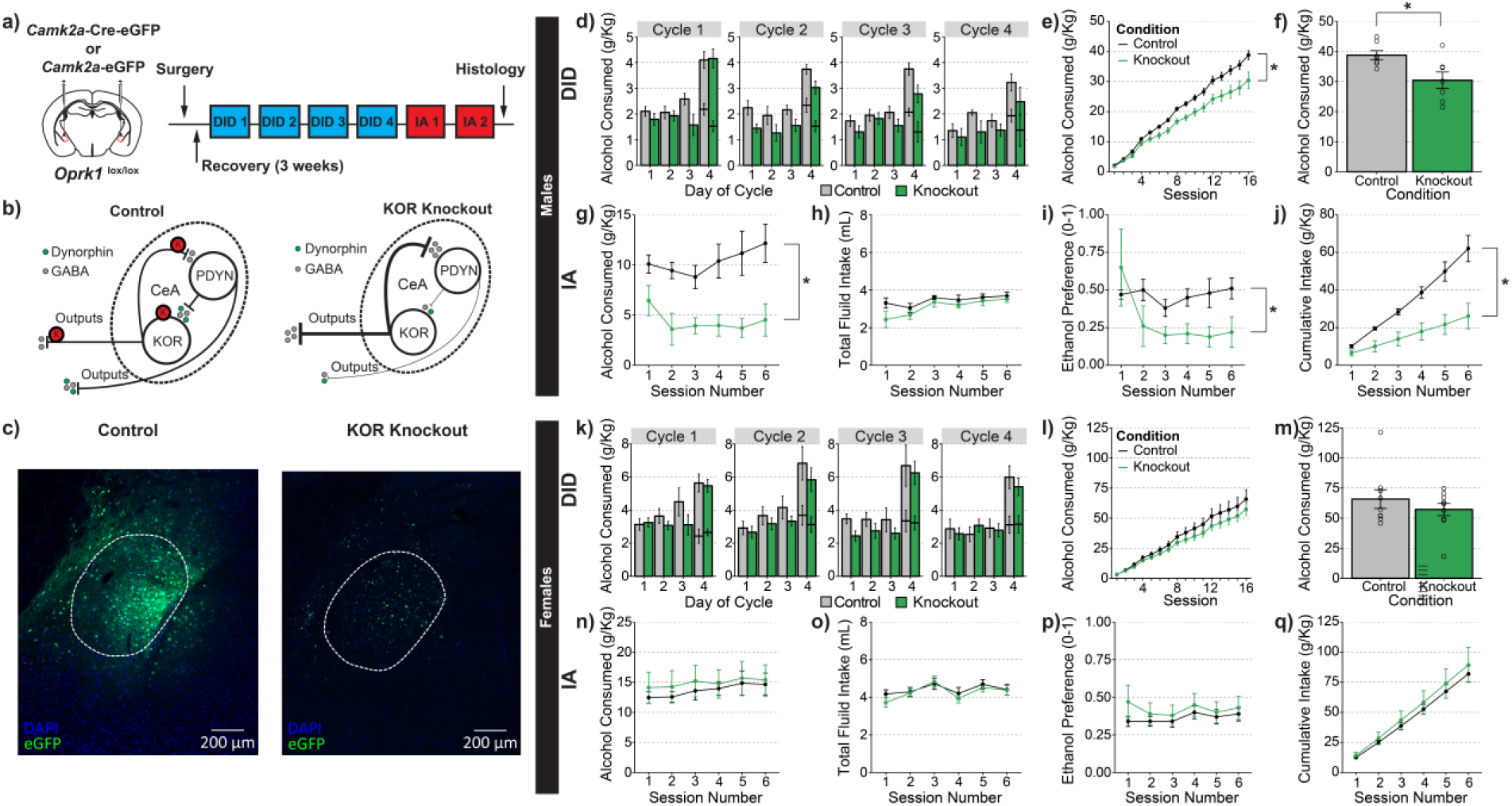
Knockout of KOR in CeA decreases ethanol consumption in male, but not female mice. (a) Timeline for experimental procedures. (b) Hypothesized actions of KOR knockout in CeA. (c) Representative images of control virus expression (left) and knockout virus (right). (d-j) Ethanol consumption in male animals. (d) Group averages for individual sessions over the course of four weeks of DID. Consumption at both the 2-hr and 4-hr time points are plotted separately on day 4. (e) Cumulative averages by group for four weeks of DID; knockout of KOR in CeA significantly decreased alcohol consumption in male animals, main effect of condition F(1,12)=7.100, p=0.021; condition X cycle interaction F(3,36)=0.324, p=0.808. (f) Cumulative totals from individual mice. (g) KOR knockout resulted in a significant reduction in the amount of alcohol consumed during IA; main effect of genotype: F(1,12)=13.628, p=0.003. (h) KOR knockout did not affect total fluid intake during IA; main effect of genotype: F(1,12)=1.652, p=0.223 (i) KOR knockout resulted in a significant reduction in ethanol preference; F(1,12)=6.616, p=0.024. (j) KOR knockout significantly reduced the cumulative amount of ethanol consumed during IA; main effect of genotype: F(1,12)=13.694, p<0.001. (k-q) Ethanol consumption in female mice. (k) Group averages for individual sessions over the course of four weeks of DID. Consumption at both the 2-hr and 4-hr time points are plotted separately on day 4. (l) Cumulative averages by group for four weeks of DID; KOR knockout in CeA did not significantly alter alcohol consumption in female animals, main effect of condition F(1,17)=0.909, p=0.353; cycle X condition interaction F(3,51)=0.614, p=0.609. (m) Cumulative totals from individual mice. (n) KOR knockout did not alter the amount of alcohol consumed during IA; main effect of genotype: F(1,17)=0.196, p=0.663. (o) KOR knockout did not affect total fluid intake during IA; main effect of genotype: F(1,17)=0.232, p=0.636. (p) KOR knockout did not alter ethanol preference; main effect of genotype: F(1,17)=0.512, p=0.484. (q) KOR knockout did not affect cumulative ethanol consumption during IA; main effect of genotype: F(1,17)= 0.263, p=0.614. (d-j) n=8 control males and n=7 knockout males (k-q) n=9 control females and n=10 knockout females

Because global KOR knockout has been shown to alter fluid intake levels (Kovacs et al., 2005), we wanted to examine whether the differences in DID observed were due to overall reductions in fluid intake. Immediately following completion of DID, experimental bottles were replaced for Intermittent Access to Ethanol (IA) so that changes in water consumption and ethanol preference could also be measured. Notably, reductions in alcohol drinking in KOR-knockout mice were observed throughout the duration of IA experiments (Figure 2g). Additionally, knockout animals had significantly lower ethanol preference, but no significant differences in the amount of total fluid consumed (Figure 2h-j).

While systemic antagonism of KOR has generally resulted in reductions in drinking in male animals (Anderson & Becker, 2017), some experiments in female animals have observed that KOR antagonism does not alter alcohol consumption (Zhou, Crowley, Ben, Prisinzano, & Kreek, 2017). Additionally, KOR activation is less aversive to female animals (Russell et al., 2014), suggesting that there may be sex differences in KOR function. Interestingly, KOR knockout in the CeA of female mice did not exhibit the reductions in alcohol drinking in DID observed in their male littermates (Figure 2k-m). Additionally, there were no significant differences in total fluid consumption or ethanol preference observed between wild-type and knockout animals during IA (Figure 2n-q). Thus, it appears that reductions in alcohol drinking demonstrated in female, global KOR-knockout animals are likely mediated outside the CeA (Kovacs et al., 2005; Van’t Veer, Smith, Cohen, Carlezon, & Bechtholt, 2016).

### Knockout of KOR in CeA Does Not Alter the Palatability of Appetitive or Aversive Tastants

One explanation for the reductions in alcohol drinking observed in the male KOR-knockout mice is that the experimental manipulation renders the taste of ethanol more aversive to knockout animals. If this were the case, the changes observed would be due to alterations in taste perception rather than the pharmacological actions of ethanol. Contrary to this hypothesis, global KOR-knockout animals are reported to have increased preference for the aversive tastant, quinine (Kovacs et al., 2005). To rigorously address this possibility, we examined a separate group of ethanol-naïve, wild-type and CeA KOR-knockout animals for their preference of a range of palatable and aversive tastant solutions (Figure S1a). We did not observe any significant differences in the quinine preference of either male or female animals for the range of concentrations tested (Figure S1b,e). Additionally, we did not observe any differences in the preferences for saccharin, a non-caloric sweetener, or sucrose, a caloric sweetener (Figure S1c,f). We also considered whether ethanol is a source of calories and the reductions in ethanol consumption in KOR-knockout males are secondary to changes in overall metabolic intake. To assess the effects of KOR knockout on the consumption of a caloric reinforcer, we performed binge access to 10% sucrose as described (Pleil, Rinker, et al., 2015; Rinker et al., 2017). We did not observe any effect of KOR knockout on binge consumption of sucrose in either male or female mice (Figure S1d,g).

### Knockout of PDYN in CeA Reduces Alcohol Consumption and Preference in a Sex-General Manner

One question arising from the previous experiment is where is the source of the endogenous dynorphin that mediates the reduction of alcohol drinking observed in male mice. Given the large adjacent population of PDYN+ neurons, one likely possibility is that it could be locally released PDYN. To selectively manipulate PDYN expression in a conditional manner, we generated homozygous *Pdyn^lox/lox^* mice to allow inactivation of *Pdyn* expression in the CeA using Cre recombinase as described in Figure S2a and Experimental Methods section. Validation with *in situ* hybridization showed that viral expression of Cre recombinase resulted in a significant reduction in the number of PDYN+ nuclei (Figure S2b-f), confirming that the *Pdyn^lox/lox^* line is an effective tool for manipulating *Pdyn* gene expression in the CeA.

To directly examine the effects of dynorphin signaling on alcohol-drinking behavior, we performed conditional knockout of PDYN in the CeA following the same timeline as that used in the KOR-knockout experiments (Figure 3a). As PDYN knockout would have the same net effect on the proposed circuit as the KOR knockout (Figure 3b), we hypothesized there would be a similar reduction in alcohol drinking by male mice. In concordance with this prediction, we observed that PDYN knockout resulted in a significant reduction in ethanol consumption across four weeks of DID in male mice (Figure 3d-f). Additionally, this reduction in alcohol drinking was observed throughout the two weeks of IA and did not significantly alter overall fluid intake (Figure 3g,h). One notable divergence from the KOR knockout is that PDYN knockout did not significantly reduce ethanol preference (Figure 3i). However, any difference may be obscured by a floor effect, as the *Pdyn^lox/+^* control animals had considerably lower preference than *Oprk1^lox/+^* control animals.

**Figure 3.**
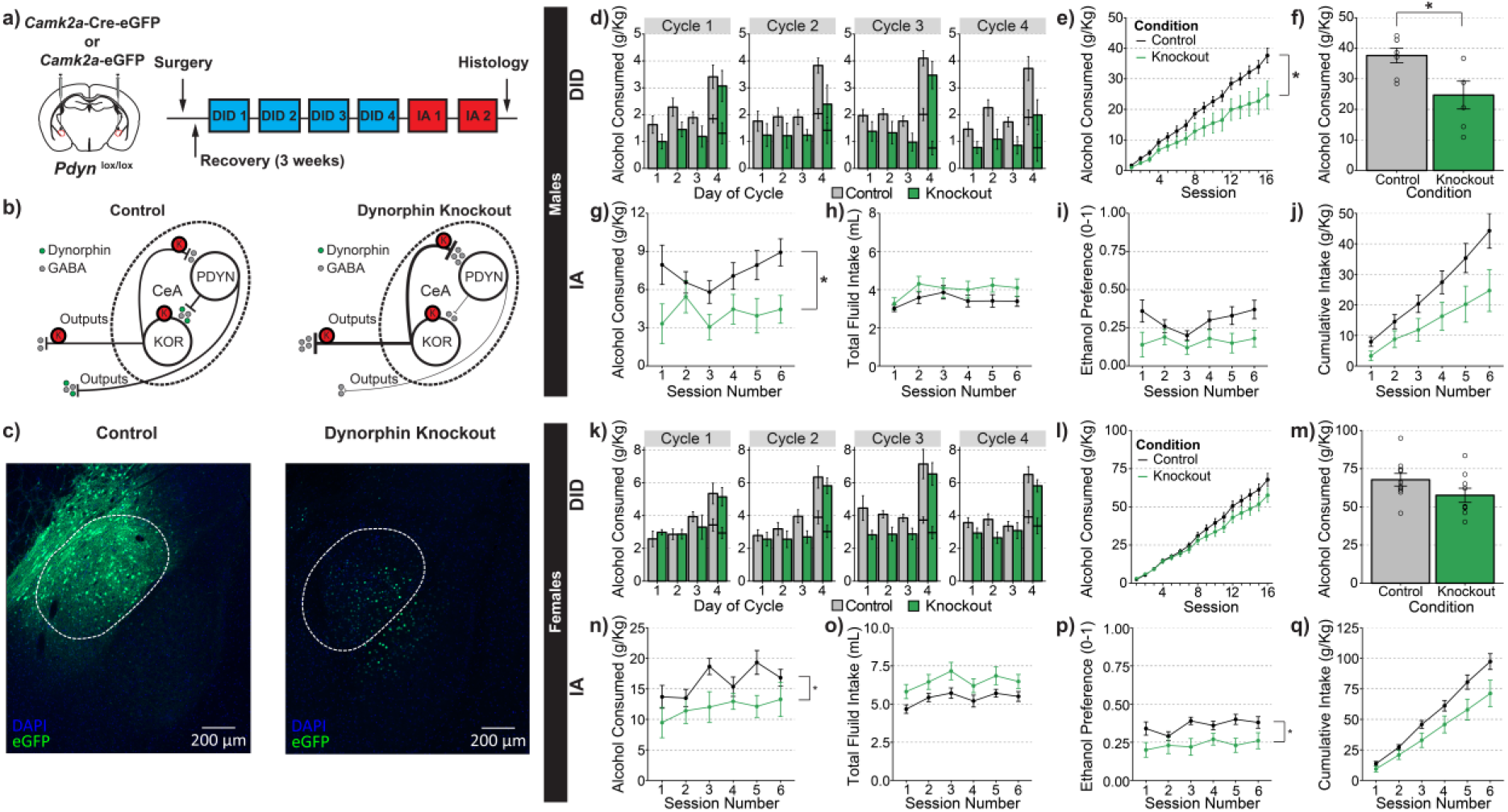
PDYN knockout in CeA decreases ethanol consumption in male and female mice. (a) Timeline for experimental procedures. (b) Hypothesized actions of PDYN knockout mice. (c) Representative images of control virus expression (left) and knockout virus (right). (d-j) Ethanol consumption in male animals. (d) Group averages for individual sessions over the course of 4 wk of DID; consumption at both 2-hr and 4-hr time points are plotted separately on day 4. (e) Cumulative averages by group for 4 wk of DID. PDYN knockout in CeA significantly decreased alcohol consumption in male animals; main effect of genotype: F(1,11)=6.899, p=0.024; genotype X cycle interaction F(3,33)=0.753, p=0.528. (f) Cumulative totals from individual mice. (g) PDYN knockout resulted in a significant reduction in the amount of alcohol consumed during IA; main effect of genotype: F(1,11)=4.860, p=0.0497. (h) PDYN knockout did not affect total fluid intake during IA; main effect of genotype: F(1,11)=1.485, p=0.248. (i) PDYN knockout did not affect ethanol preference; main effect of genotype: F(1,11)=3.788, p=0.078. (j) PDYN knockout did not significantly reduce the cumulative amount of ethanol consumed during IA F(1,11)=3.943, p=0.073. (k-q) Ethanol consumption in female animals. (k) Group averages for individual sessions over the course of 4 wk of DID. Consumption at both the 2-hr and 4-hr points are plotted separately on day 4. (l) Cumulative averages by group for 4 wk of DID; knockout of PDYN in CeA did not significantly alter alcohol consumption in female animals; main effect of genotype: F(1,17)=2.707, p=0.118; genotype X cycle interaction F(3,51)=1.643, p=0.191. (m) Cumulative totals from individual mice. (n) PDYN knockout significantly reduced the amount of alcohol consumed during IA; main effect of genotype: F(1,17)=4.490, p=0.049. (o) PDYN knockout did not affect total fluid intake during IA; main effect of genotype: F(1,17)=4.450, p=0.501. (p) PDYN knockout significantly reduced ethanol preference; main effect of genotype: F(1,17)=6.766, p=0.019. (q) PDYN knockout did not affect cumulative ethanol consumption during IA; main effect of genotype: F(1,17)=4.135, p=0.579 (d-j) n=7 control males and n=6 knockout males. (k-q) n=10 control females and n=9 knockout females

We also tested whether knockout of PDYN in CeA would result in similar reductions in female mice, or if there would be a divergent response between sexes as was observed with KOR knockout. Thus, we deleted PDYN in CeA of female mice following the same timeline as before. We observed that CeA PDYN lowered, but did not significantly reduce, ethanol consumption during DID (Figure 3k-m). However, PDYN knockout did significantly reduce alcohol consumption during IA in females (Figure 3n) and reduced ethanol preference without affecting overall fluid intake (Figure 3o,p). Thus, it appears that PDYN knockout in the CeA does play some role in regulating ethanol consumption in female mice.

### PDYN Knockout in CeA Does Not Alter the Palatability of Appetitive or Aversive Tastants

It has been demonstrated that global PDYN-knockout mice have significantly altered fluid intake, saccharin preference, quinine preference, and sucrose consumption compared to littermate controls (Blednov et al., 2006). To assess the effects of CeA PDYN knockout on each of these parameters, ethanol-naïve mice underwent tastant drinking experiments following the same timeline as KOR-knockout experiments (Figure S3a). In contrast to what was observed in the PDYN global-knockout animals, PDYN knockout in CeA of male mice did not alter quinine, saccharin, or sucrose preference (Figure S3 b,c). Additionally, PDYN knockout did not alter binge consumption of a 10% sucrose solution (Figure S3d). Similarly, PDYN knockout in female mice did not have an effect on the preference of any of the tastant solutions (Figure S3e,f) or limited access consumption of sucrose (Figure S3g). Thus, it is likely that PDYN knockout in CeA selectively alters the drive to consume alcohol without affecting general taste perception.

### Neither KOR Nor PDYN Knockout Protects Mice Against Alcohol Drinking-Induced Increases in Anxiety

Ethanol dependence has been shown to increase anxiety-like behavior in the elevated-plus maze (EPM), which can be reversed by systemic treatment with a KOR antagonist (Valdez & Harshberger, 2012). If PDYN or KOR knockout is protective against the negative reinforcing properties of ethanol, these manipulations should produce a detectable change in anxiety-like behavior during acute withdrawal. Because CeA KOR knockout decreased ethanol consumption, we examined the effects of CeA KOR knockout in both ethanol-naïve animals and those that underwent DID (Figure 4a). We observed that there was a main effect of alcohol drinking whereby a history of alcohol use increased anxiety-like behavior in the EPM in male mice (Figure 4bc). However, there was no main effect or interaction with genotype, indicating that KOR knockout does not alter anxiety in the basal state or during withdrawal. Similar effects were observed in female mice, as there was a main effect of alcohol drinking on open-arm entries, but no main effect or interaction with genotype (Figure 4f). With PDYN-knockout mice, we did not observe any effects of ethanol drinking on open-arm time or entries (Figure 4 h-i). However, ethanol drinkers had significantly higher locomotion than naïve controls (Figure 4j). In female mice, there was no effect of PDYN knockout on ethanol drinking, open-arm time, open-arm entries, or total distance traveled (Figure 4 k-m).

**Figure 4.**
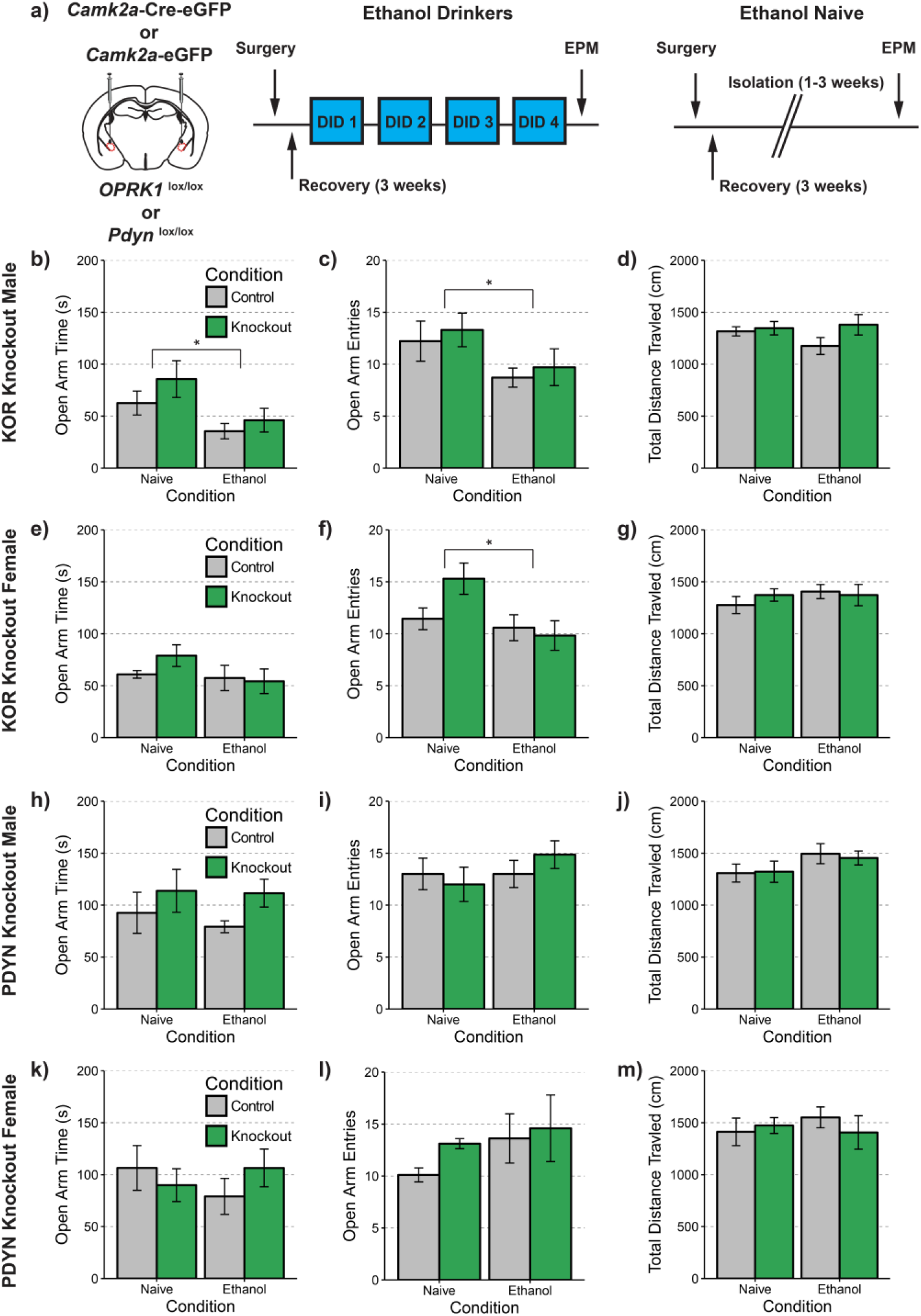
Neither KOR nor PDYN Knockout Protect Against Ethanol-Induced Increases In Anxiety-Like Behavior. (a) Experimental timeline. (b) There was a significant main effect of ethanol drinking F(1,29)=5.739, p=0.023 but not gene knockout F(1,29)=1.651, p=0.209 on open arm time. (c) There was a significant main effect of ethanol drinking F(1,29)=4.372, p=0.045 but not gene knockout F(1,29)=0.388, p=0.538 on open arm entries. (d) There was no effect of ethanol drinking F(1,29)=0.522, p=0.476 or gene knockout F(1,29)=2.18, p=0.151 on total distance traveled in the elevated plus maze. (e) There was no significant effect of ethanol drinking F(1,39)= 1.765, p=0.192 or gene knockout F(1,39)=0.332, p=0.568 on open arm time. (f) There was a significant main effect of ethanol drinking F(1,39)=5.718, p=0.022 but not gene knockout on open arm entries. (g) There was no effect of ethanol drinking F(1,39)=0.582, p=0.450 or gene knockout F(1,39)=0.081, p=0.777 on total distance traveled. (h) There was no effect of ethanol drinking F(1,29)=0.197, p=0.660 or gene knockout F(1,29)=2.149, p=0.153 on open-arm time. (i) There was no effect of ethanol drinking F(1,29)=0.827, p=0.371 or gene knockout F(1,29)=0.020, p=0.889 on open-arm entries. (j) There was no significant main effect of ethanol treatment F(1,29)=3.009, p=0.093 or gene knockout F(1,29)=0.011, p=0.915 on total distance traveled. (k) There was no effect of ethanol drinking F(1,26)=0.213, p=0.648, or gene knockout F(1,26)=0.009, p=0.922 on open-arm time. (l) There was no effect of ethanol drinking F(1,26)=2.374, p=0.135 or gene knockout F(1,26)=1.564, p=0.222 on open-arm entries. (m) There was no effect of ethanol drinking F(1, 26)=0.198, p=0.660 or gene knockout F(1, 26)=0.048, p=0.829 on total distance traveled. (b-d) n=8 ethanol control males, n=7 ethanol knockout males, n=9 naïve control males, n=7 naïve knockout males; (e-g) n=9 ethanol control females, n=10 knockout females, n=7 naïve control females, n=10 knockout females; (h-j) n=7 ethanol control males, n=7 ethanol knockout males, n=10 naïve control males, n=9 naïve knockout males; (k-m) n=8 ethanol control females, n=7 knockout females, n=9 naïve control females, n=8 naïve knockout females.

### A History of Ethanol Drinking Does Not Alter Synaptic Transmission onto PDYN Neurons in the CeA

A primary role ascribed to activation of KOR signaling in the CeA is that it reduces presynaptic GABA release (Gilpin, Roberto, Koob, & Schweitzer, 2014; Kang-Park, Kieffer, Roberts, Siggins, & Moore, 2013). To assess whether KOR modulation of inhibitory synaptic transmission was altered after a history of binge-like alcohol drinking, we performed slice-electrophysiology recordings in slices from the CeA of *Pdyn^IRES-Cre^* :: *Gt(ROSA26)Sor^loxSTOPlox-L10-GFP^* mice that have fluorescent ribosomes only in PDYN neurons (see Experimental Methods). The mice underwent three cycles of DID with access to 20% ethanol or water (Figure 5a). As reported previously, we found that the KOR agonist U-69593 (1 μM) reduced electrically evoked inhibitory post synaptic current (eIPSC) amplitude in male mice (Figure 5b). However, the degree of inhibition was unaltered when comparing ethanol and water drinkers. We next examined the effect of KOR activation on eIPSC amplitude in female PDYN-reporter mice, and found that KOR inhibition of GABAergic transmission was similarly unaltered by a history of alcohol drinking in female mice.

**Figure 5.**
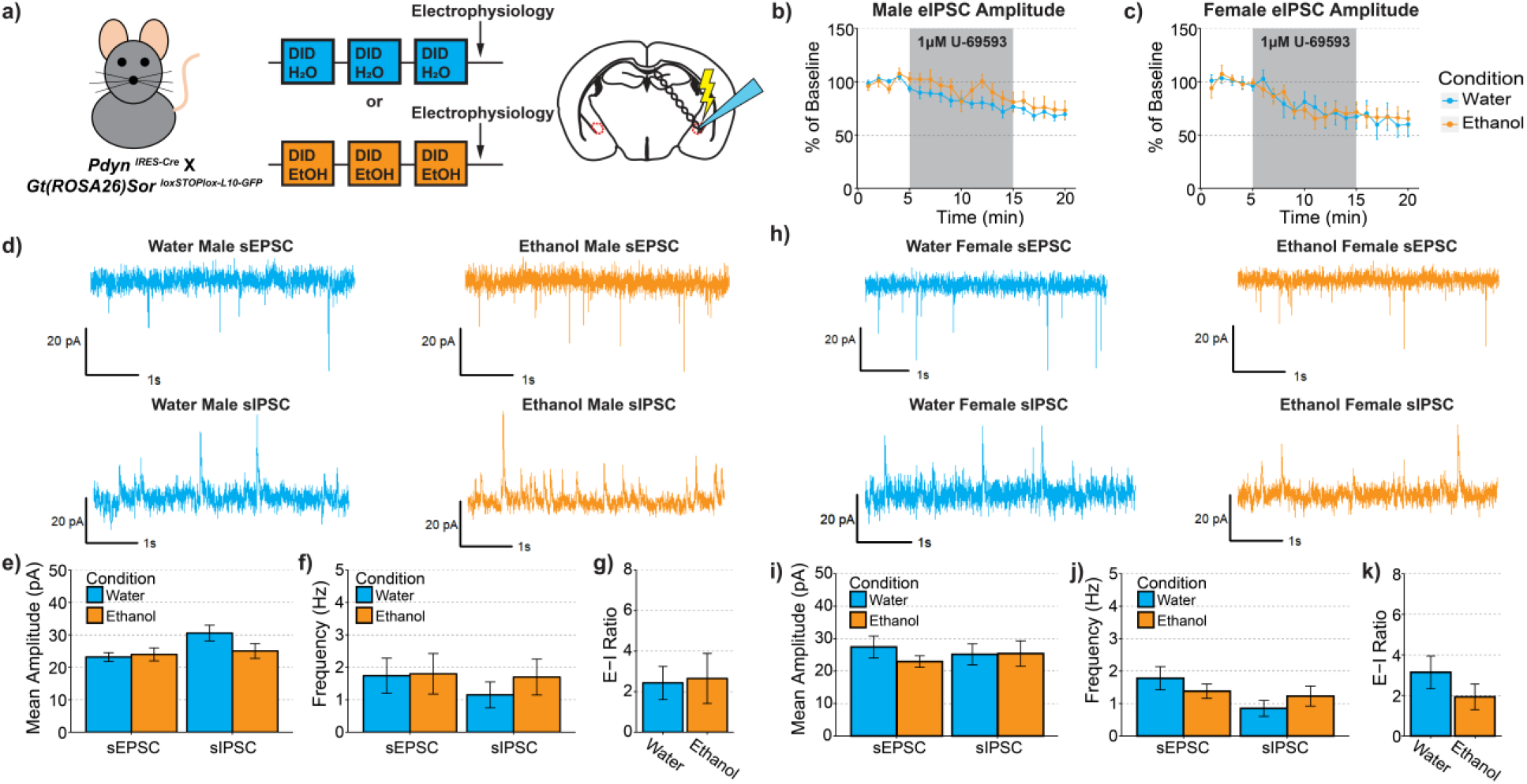
Ethanol Drinking Does Not Alter Synaptic Transmission Onto CeA PDYN Neurons. (a) Experimental timeline. (b) U69593 inhibition of eIPSCs is unaltered by a history of alcohol drinking in males t(9.391)=0.687, p=0.508. (c) U69593 inhibition of eIPSCs is also unaltered by a history of alcohol drinking in females t(14.880)=0.054, p=0.957. (d-g) Synaptic transmission in male animals. (d) Representative traces of excitatory and inhibitory events onto CeA dynorphin neurons. (e) Ethanol drinking did not alter the amplitude of excitatory t(11.152)=0.334, p=0.745 or inhibitory transmission t(14.704)=1.644, p=0.121. (f) Ethanol drinking did not alter the frequency of excitatory transmission t(13.390)=0.075, p=0.941 or inhibitory transmission t(11.767)=0.815, p=0.431. (g) synaptic drive was also unaltered after alcohol drinking t(10.982)=0.153, p=0.881. (h-k) Synaptic transmission in female animals. (h) Representative traces of excitatory and inhibitory events onto CeA dynorphin neurons. (i) Ethanol drinking did not alter the amplitude of excitatory transmission t(9.164)=1.170, p=0.272 or inhibitory transmission t(12.847)=0.046, p=0.964. (j) Ethanol drinking did not alter the frequency of excitatory transmission t(10.198)=0.966, p=0.356 or inhibitory transmission t(12.731)=0.950, p=0.359. (k) synaptic drive was also unaltered after alcohol drinking t(11.806)=1.182, p=0.260. (b) Water male n=9 cells, N=6 mice, ethanol male n=7 cells, n=4 mice; (c) water female n=7 cells, N=5 mice, ethanol female n=6 cells, n=4 mice; (d-g) water male n=10 cells, n=6 mice, ethanol male n=9 cells, n=4 mice; (h-k) water female n=7 cells, n=4 mice, ethanol female n=8 cells, n=5 mice

It has been shown that a history of alcohol exposure can alter the balance of excitatory and inhibitory transmission in the CeA (Pleil, Lowery-Gionta, et al., 2015; Roberto, Madamba, Stouffer, Parsons, & Siggins, 2004; Roberto, Schweitzer, et al., 2004). Thus, we assessed whether binge-like alcohol drinking alters spontaneous synaptic transmission onto PDYN neurons in the CeA. In male mice, we did not observe any effect of alcohol drinking on either the frequency, amplitude or decay kinetics of excitatory or inhibitory transmission (Figure 5 d-g; Table 2 & 3). Spontaneous synaptic transmission in female mice was also unaffected by alcohol drinking (Figure 5 h-k). These experiments suggest that the protective effects of PDYN/KOR knockout are likely not mediated by altering the balance of excitatory and inhibitory transmission in the CeA.

**Table 1.** Cell Properties in Synaptic-Transmission Experiments

|  | Water |  | Ethanol |  |
| --- | --- | --- | --- | --- |
| | Capacitance (pF)<br>± SEM | Resistance (MΩ)<br>± SEM | Capacitance (pF)<br>± SEM | Resistance (MΩ)<br>± SEM |
| Male | 77.11 ± 7.58 | 313.96 ± 73.19 | 85.32 ± 6.45 | 259.26 ± 35.44 |
| Female | 81.26 ± 6.68 | 388.67 ± 79.45 | 82.38 ± 6.52 | 210.51 ± 19.44 |

**Table 2.** Kinetics for spontaneous Excitatory Post-synaptic Currents (sEPSCs)

|  | Water |  | Ethanol |  |
| --- | --- | --- | --- | --- |
| | Rise (ms) ± SEM | Decay (ms) ± SEM | Rise (ms) ± SEM | Decay (ms) ± SEM |
| Male | 1.91 ± 0.14 | 3.79 ± 0.16 | 2.01 ± 0.11 | 5.17 ± 0.89 |
| Female | 1.90 ± 0.20 | 3.62 ± 0.44 | 2.14 ± 0.22 | 4.74 ± 0.72 |

**Table 3.** Kinetics for spontaneous Inhibitory Post-synaptic Currents (sIPSCs)

|  | Water |  | Ethanol |  |
| --- | --- | --- | --- | --- |
| | Rise (ms) ± SEM | Decay (ms) ± SEM | Rise (ms) ± SEM | Decay (ms) ± SEM |
| Male | 3.79 ± 0.28 | 13.01 ± 1.03 | 4.17 ± 0.53 | 10.55 ± 1.41 |
| Female | 3.75 ± 0.26 | 10.61 ± 1.38 | 4.05 ± 0.37 | 12.20 ± 1.54 |

**Table 4.** Cell Properties in Excitability Experiments

|  | Water |  | Ethanol |  |
| --- | --- | --- | --- | --- |
| | Capacitance (pF)<br>± SEM | Resistance (MΩ)<br>± SEM | Capacitance (pF)<br>± SEM | Resistance (MΩ)<br>± SEM |
| Male | 74.33 ± 9.00 | 139.44 ± 24.24 | 65.81 ± 8.24 | 156.27 ± 16.32 |
| Female | 68.93 ± 7.39 | 222.63 ± 36.61 | 61.71 ± 7.29 | 163.58 ± 46.36 |

### Ethanol Drinking Results in Sex-Specific Changes in Intrinsic Excitability of PDYN Neurons in the CeA

Alcohol exposure has been shown to result in cell-type specific changes in the excitability of CeA neurons (Herman, Contet, & Roberto, 2016). To assess whether alcohol drinking may alter cell intrinsic excitability, we examined whether a history of alcohol drinking altered the firing properties of PDYN neurons in CeA. In these experiments, the reporter mice underwent DID with 20% ethanol or water as described above. In male mice, ethanol drinking did not alter the resting membrane potential, action-potential threshold, or rheobase of PDYN neurons (Figure 6a-c). However, ethanol drinkers fired significantly more action potentials in response to increasing steps of depolarizing current (Figure 6d-e). In female mice, resting membrane potential, action-potential threshold, and rheobase were similarly unaltered after alcohol exposure (Figure 6f-h). There was also a small, but insignificant trend towards reduced number of action potentials fired in a voltage versus current plot (Figure 6j). These experiments demonstrate that a history of alcohol drinking changes the excitability of PDYN neurons in opposing directions in male and female mice, which may partially explain the divergent results seen with manipulations *in vivo*.

**Figure 6.**
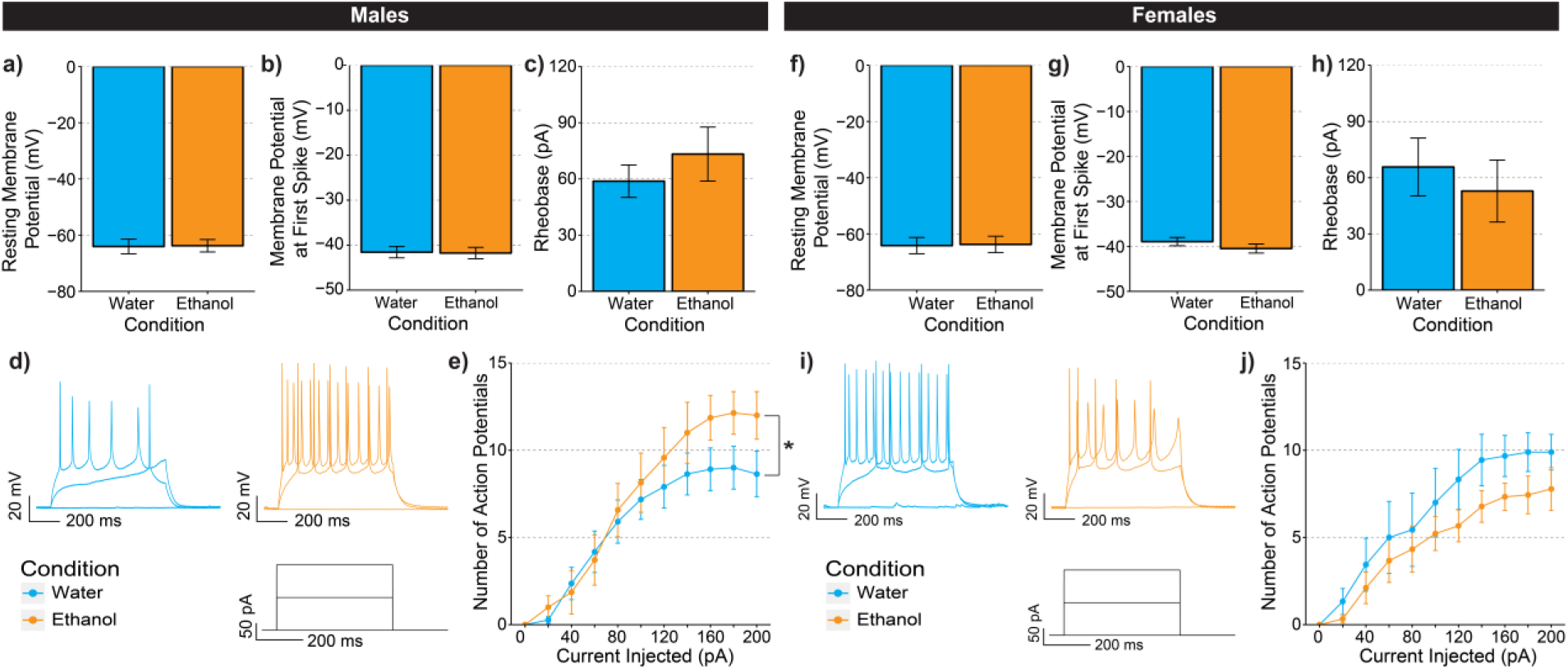
Ethanol Drinking Alters the Excitability of PDYN Neurons in A Sex-Specific Manner. (a-e) Data collected from male mice. (a)There was no effect of ethanol drinking on resting membrane potential t(16.833)=0.076, p=0.940. (b) Ethanol drinking did not change action potential threshold t(16.947)=0.111, p=0.912. (c) There was no significant difference in the amount of current injected needed to elicit an action potential t(13.279)=0.860, p=0.405. (d) Representative traces of PDYN-neuron firing in CeA. (e) There was a significant interaction between ethanol drinking and the number of action potentials fired in the VI plot F(10,160)=2.004, p=0.036. (f-j) Data collected from female mice. (f) There was no effect of ethanol drinking on resting membrane potential t(13.681)=0.101, p=0.920. (g) Action potential threshold t(13.248)=1.120, p=0.283 or (h) rheobase t(13.386)=0.568, p=0.579. (i) Representative traces of CeA PDYN-neuron firing. (j) Ethanol drinking resulted in a nonsignificant decrease in the number of action potentials fired in the VI plot F(1,16)=2.523, p=0.132. (a-e) Water males: n=10 cells from n=6 mice; ethanol male: n=9 cells from N=6 mice; (f-j) water females: n=8 cells from n=5 mice; ethanol female n=9 from n=5 mice

## Discussion

Despite its high prevalence, there remain only three FDA approved pharmacotherapies available to treat AUD (Heilig & Egli, 2006). One of these therapies, naltrexone, works as a non-specific opioid antagonist. Recently, there has been an interest in developing KOR-specific antagonists to treat AUD as well as other psychiatric conditions (Carroll & Carlezon, 2013; Chavkin & Koob, 2016; Crowley & Kash, 2015). Understanding how KOR signaling works in the amygdala is critically important with respect to AUD, as a recent study found reduced KOR occupancy in the amygdala in alcoholics, consistent with tonic *in vivo* engagement of DYN/KOR signaling (Vijay et al., 2018). The present study sought to address this by mechanistically dissecting how PDYN and KOR signaling in the CeA contribute to excessive alcohol intake. We observed that PDYN and KOR form spatially adjacent, but mostly distinct populations suggesting that the balance of these two cell types may control the balance of different outputs. We then performed separate knockouts of PDYN and KOR and found that while knockout of PDYN decreased alcohol consumption in both sexes, knockout of KOR resulted in reductions of alcohol intake in male mice only. Finally, we found sex-specific reductions in PDYN neuron excitability in female mice following alcohol drinking.

There are several important nuances to our findings. First, the mechanism of protective action of PDYN or KOR knockout is likely different between sexes. The fact that KOR knockout in CeA did not affect alcohol drinking in female mice may not be surprising given that KOR activation has been shown to be less aversive to female animals (Russell et al., 2014). However, it should be noted that female KOR global knockout animals showed reduced ethanol consumption and preference comparable to their male counterparts (Kovacs et al., 2005). In the present study, the results in female mice were not entirely negative; PDYN knockout produced a significant reduction in ethanol consumption and preference in IA. Thus, our findings are consistent with a model in which PDYN neurons may promote alcohol consumption in females, but these effects may be mediated independently of KOR in CeA neurons. One alternative is that PDYN neurons could be sending projections to other brain regions that express KOR, including the BNST, and terminal release of dynorphin in these regions could drive changes in behavior independent of KOR expressed in the CeA. Another possibility is that the effects of PDYN knockout in females may be mediated by opioid peptides other than dynorphin. As *Pdyn* also encodes leu-enkephalin and β-neoendorphin, the reductions in alcohol drinking seen in female CeA PDYN knockout mice may instead be mediated by one of these other peptides.

The two sexes were also divergent at the level of CeA PDYN neuron plasticity. We observed that a history of ethanol drinking increased the excitability of PDYN neurons in male mice, but exhibited a trend towards decreased excitability in female mice. In the present study, we did not observe any effects of DID on spontaneous excitatory and inhibitory transmission in the CeA of either sex. Consistent with previous results, we found that KOR activation reduces eIPSC amplitude in male mice and demonstrates for the first time that KOR activation similarly decreases eIPSC amplitude in female mice as well. However, we did not find any effects of ethanol drinking on KOR inhibition of GABAergic transmission in either sex. Future studies should examine whether a history of binge-like drinking results in alterations in endogenous PDYN tone.

Another important finding was that neither PDYN nor KOR knockout did not protect against ethanol withdrawal-induced increases in anxiety. While ethanol vapor exposure and ethanol liquid diet have been shown to increase anxiety-like behavior in rodent models (Overstreet, Knapp, & Breese, 2004; Pleil, Lowery-Gionta, et al., 2015; Rose et al., 2016; Valdez & Harshberger, 2012), increased anxiety after DID typically has not been observed (Cox et al., 2013). However, these studies assessed anxiety using the EPM and open-field paradigms. Lee et al. (2018) have shown that increased anxiety-like behavior can be observed after DID in the light-dark assay suggesting that changes in anxiety may not be revealed by all assays. Another possibility is that the differences observed may be due to the timing of when the testing took place relative to the last ethanol exposure session. We performed EPM testing 8 hr after the last binge session based on the finding that withdrawal symptoms peak around 6-10 hr after ethanol (Becker & Hale, 1993). Because testing in the study by Cox et al. (2013) was conducted 24 hr after the last binge session, it is possible that their experiments may have missed transient changes in anxiety like behavior. Additionally, there was no significant main effect of PDYN or KOR knockout or interaction with ethanol treatment on anxiety-like behavior suggesting that the reductions in alcohol drinking were mediated without concomitant changes in anxiety levels.

One limitation of our study is that we did not examine the outputs of CeA PDYN and KOR neurons. It has been demonstrated that optogenetic inhibition of CRF-expressing neurons in the CeA that project to the BNST reduces ethanol vapor-induced increases in ethanol self-administration (Guglielmo et al., 2019). Given the high overlap of PDYN and CRF expression in the CeA (Kim et al., 2017), PDYN projections to the BNST are a likely output that mediates these effects. Additionally, because we hypothesize the balance of excitation between these two populations is important for regulating alcohol intake, it would be informative to trace the outputs of CeA KOR neurons. Identifying any common or divergent outputs of these two populations would provide further insight into what behavioral processes may be affected. As *Oprk1* is also expressed in the BLA, our injections may have resulted in a small degree of KOR knockout in the BLA. However, we did not observe the BLA KOR-mediated reductions in anxiety in these subjects (Figure 4b) suggesting that these effects are mediated by CeA KORs.

The major advancement of the present study was to identify the effects of PDYN or KOR signaling in the CeA in regulating alcohol drinking. Additionally, they do so in a sex-specific manner without affecting general taste preference or anxiety-like behavior. These findings contribute to our growing appreciation that the specific effects of KOR signaling in different brain regions may diverge from effects seen with global manipulations. Indeed, PDYN or KOR activation in the NAc has led to both appetitive and aversive responses based on anatomical sub-region that was targeted (Al-Hasani et al., 2015; Castro & Berridge, 2014). Additionally, as KOR may be differentially expressed on different cell types within a brain region and shape output (Tejeda et al., 2017), it is important to examine the effects of KOR activation in the projection-specific circuits. For example, KOR inhibition of GABAergic projections to the BNST are mediated through ERK1/2, whereas KOR inhibition of glutamatergic BLA inputs are mediated by p38/MAP Kinase (Crowley et al., 2016; Li et al., 2012). These differences can then be exploited therapeutically with biased agonists (Bohn & Aubé, 2017; Brust et al., 2016; Ho et al., 2018) and could then be used to shift the balance of inputs from one brain region to another.

Taken together, our findings support the continued search for KOR therapeutics to treat AUD. Moreover, they suggest that KOR antagonism as a treatment modality may exhibit sex differences if excessive alcohol consumption is driven via amygdala KOR signaling.

## Acknowledgments

We would like to thank Christina Catavero and Christina Stanhope for assistance with histology and alcohol drinking measurements. We would also like to thank Maria Luisa Torruella-Suarez and Zoe McElligott for thoughtful discussion.

## Conflict of Interests

The authors have no conflict of interests to disclose.

---

**Supplementary Figure 1.**
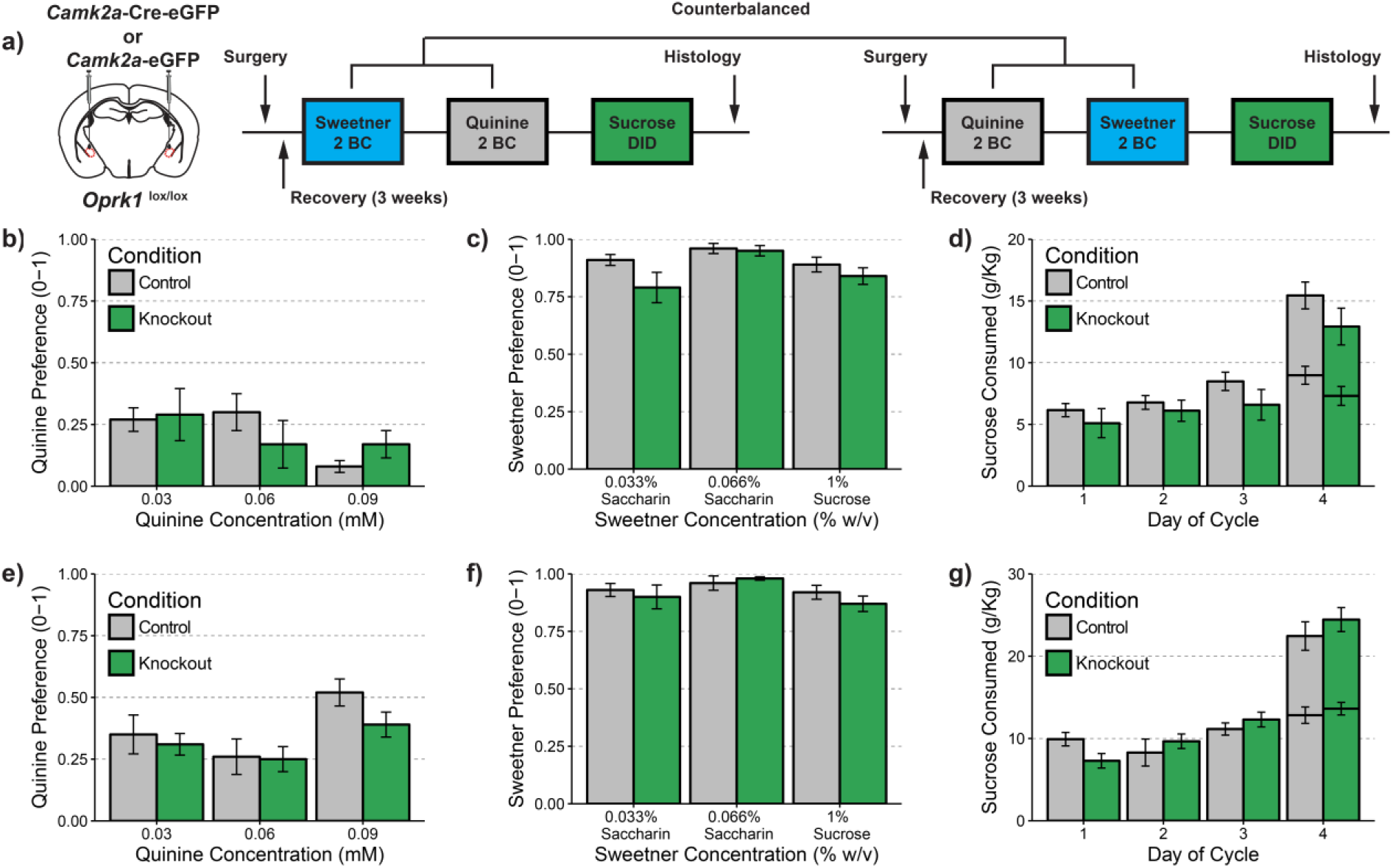
CeA KOR knockout does not alter the preference of palatable or aversive tastants. (a) Timeline for tastant experiments. Ethanol-naïve animals underwent two-bottle choice (2BC) for series of palatable (saccharin and sucrose) or aversive (quinine) tastants. The presentation of the first series (palatable or aversive) was counterbalanced across multiple cohorts. Following completion of the two-bottle choice experiments, animals then had binge access to 10% sucrose during same access schedule as DID. (b-d) Experiments in male animals. (b) CeA KOR knockout did not alter preference for quinine at the range of concentrations tested, main effect of genotype; F(1,14)=0.005, p=0.945. (c) KOR knockout did not alter for preference for saccharin or sucrose, main effect of genotype; F(1,14)=3.015, p=0.104. (d) KOR knockout did not affect binge consumption of sucrose at any of the 2-hr time points; main effect of genotype: F(1,12)=1.435, p=0.254 or 4-hr time point on the final day t(13.168), p=0.193. (e-g) Experiments in female animals. (e) CeA KOR knockout did not alter preference for quinine at the range of concentrations tested, main effect of genotype: F(1,14)=2.069, p=0.172. (f) KOR knockout did not alter for preference for saccharin or sucrose, main effect of genotype: F(1,14)=0.175, p=0.682. (g) KOR knockout did not affect binge consumption of sucrose at any of the 2-hr time points, main effect of genotype: F(1,14)=0.007, p=0.979, or 4-hr time point on the final day t(11.655), p=0.397. (b-d) n=9 control males and n=7 knockout males, (e-g) n=7 control females and n=10 knockout females

**Supplementary Figure 2.**
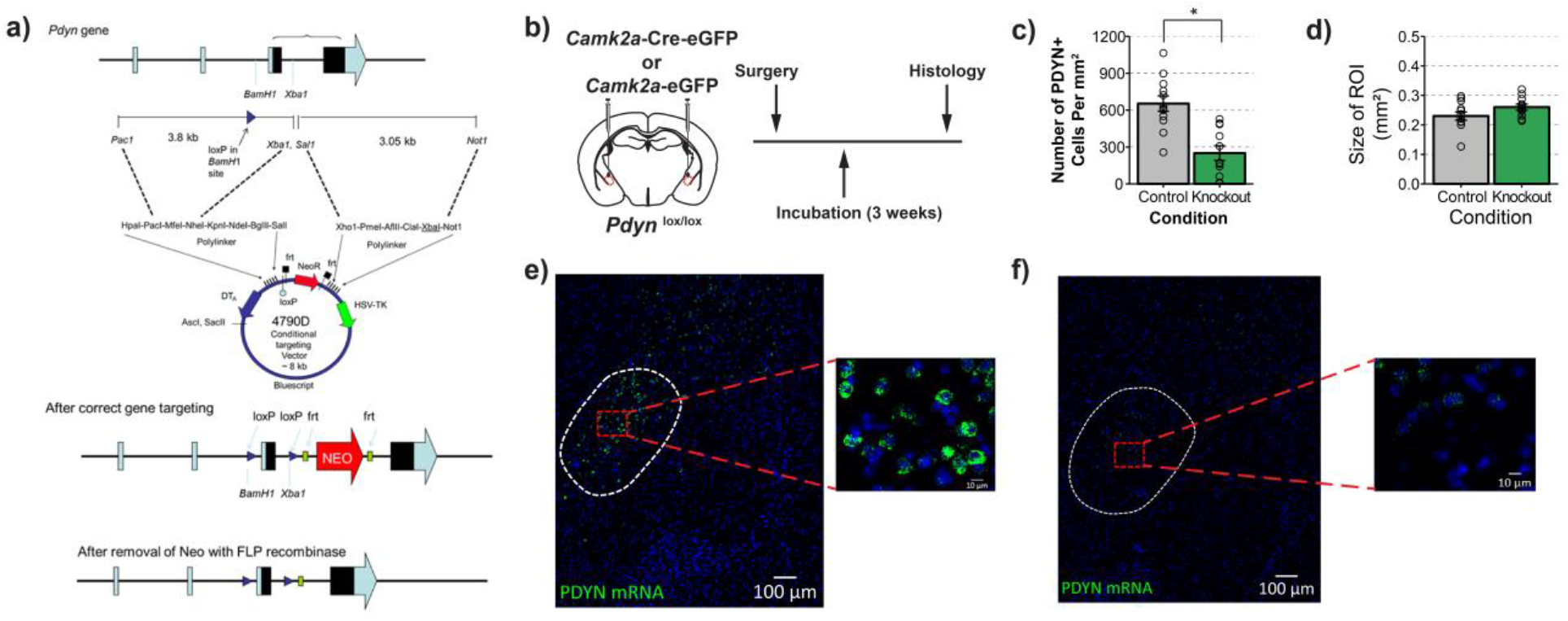
Validation of PDYN-Flox Line. (a) Cloning vector used to target *Pdyn* gene. (b) Experimental timeline for virus injection and histology. (c) Injection of an adeno associated virus containing Cre recombinase results in a significant reduction in the number of PDYN+ cells in the CeA t(21)= 4.663, p<0.001. (d) The size of the ROI drawn was no different between groups t(19.953)=1.660, p=0.113. (e) Representative image of *Pdyn* mRNA expression in a control mouse and (f) in a PDYN-knockout mouse. n=12 images per group from n=4 mice per condition.

**Supplementary Figure 3.**
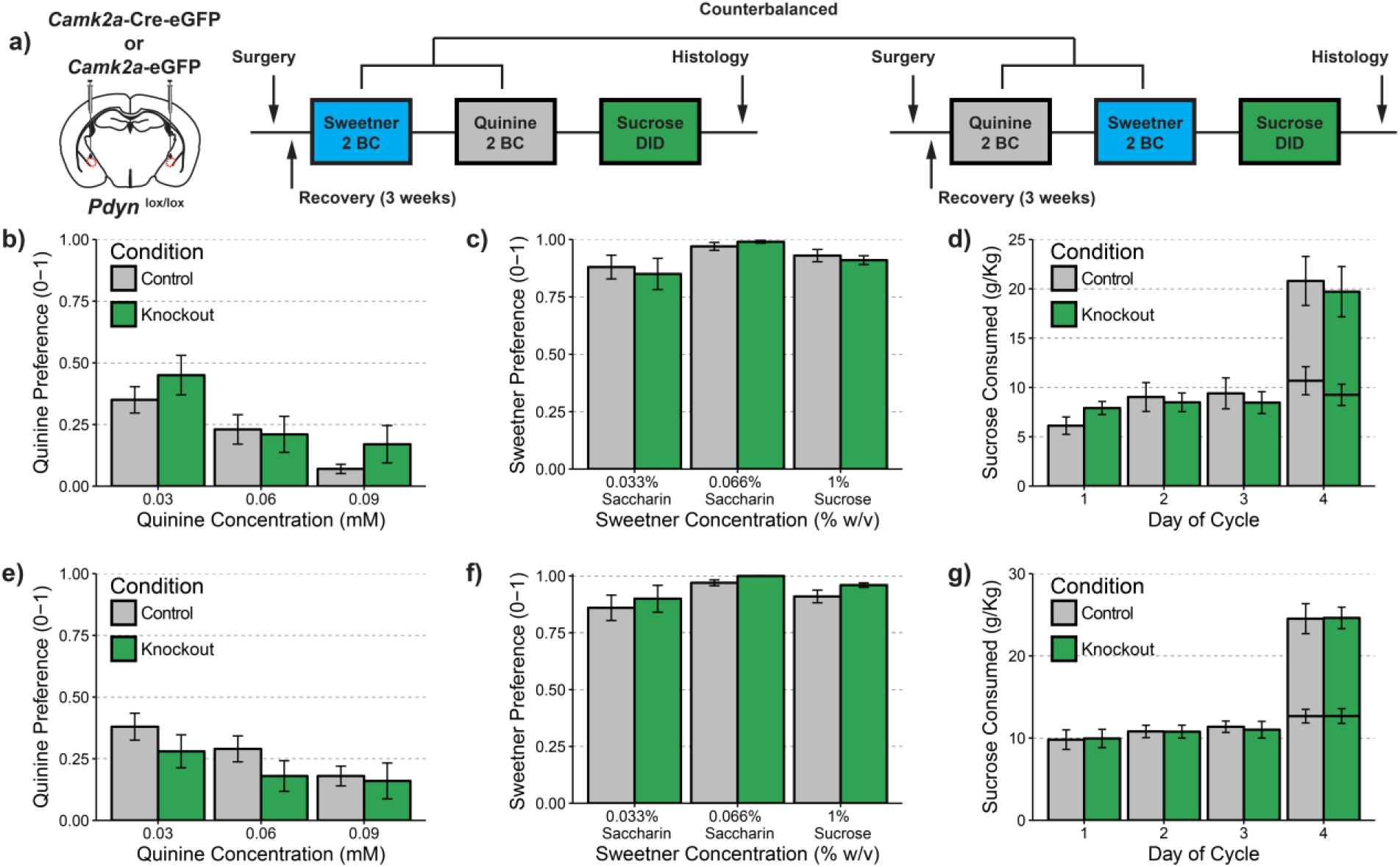
PDYN knockout in CeA does not alter the preference of palatable or aversive tastants. (a) Timeline for tastant experiments. Ethanol naïve animals underwent two-bottle choice (2BC) for series of palatable (saccharin and sucrose) or aversive (quinine) tastants. The presentation of the first series (palatable or aversive) was counterbalanced across multiple cohorts. Following completion of the two-bottle choice experiments, animals then had binge access to 10% sucrose during same access schedule as DID. (b-d) Experiments in male animals. (b) CeA PDYN knockout did not alter preference for quinine at the range of concentrations tested; main effect of genotype: F(1,13)=0.752, p=0.402. (c) PDYN knockout did not alter for preference for saccharin or sucrose, main effect of genotype: F(1,13)=0.093, p=0.765. (d) PDYN knockout did not affect binge consumption of sucrose at any of the 2-hr time points; main effect of genotype: F(1,11)=0.322, p=0.582 or 4-hr time point on the final day t(12.963)=0.306, p=0.764. (e-g) Experiments in female mice. (e) CeA PDYN knockout did not alter preference for quinine at the range of concentrations tested, main effect of genotype: F(1,13)=1.119, p=0.309. (f) PDYN knockout did not alter for preference for saccharin or sucrose, main effect of genotype: F(1,13)=1.331, p=0.269. (g) PDYN knockout did not affect binge consumption of sucrose at any of the 2-hr time points, main effect of genotype: F(1,12)=1.184, p=0.298 or 4-hr time point on the final day t(13.18)=0.325, p=0.750. (b-d) n=7 control males and n=8 knockout males (e-g) n=8 control females and n=7 knockout females.

